# Profiling susceptibility of NIAID Category A and B priority and emerging pathogens to define strain panels for drug discovery and active drug classes

**DOI:** 10.1101/2020.12.04.412767

**Authors:** Jason E. Cummings, Zaid Abdo, Richard A. Slayden

## Abstract

Drug susceptibility profiles of NIAID Category A and B priority and emerging pathogens to standard of care drugs were assessed to determine susceptibility against clinical strains in addition to reference laboratory strains to establish a comparison resource for the performance of drug candidates, to define non-redundant minimal strain panels for individual species and the group of species for use in drug screening programs, and provide pharmacophore classification. Profiling of standard of care drugs against strains of each species revealed a broad spectrum of susceptibility among strains in each species with important differences between standard laboratory reference strains and strains sof clinical origin. Unbiased hierarchical clustering analyses of strain susceptibilities within each species group and strains from all the species identified subsets of non-redundant strains that are able to classify the susceptibility range for each species and able to classify all the species. This analysis established a reduced targeted set of Category A and B priority pathogen strains for testing the potency of drug candidates against each species and pan-species. This approach also discriminated pharmacophore classification for each species. This information can be applied to directed screening efforts to guide the selection of drug classes for derivatization and repurposing, and advance drug candidates with the greatest potential for efficacy against NIAID Category A and B priority and emerging pathogens.

## Introduction

NIAID Category A and B priority pathogens are included in the classification of emerging infectious diseases that have been defined as those caused by organisms that are newly appearing or increasing in incidence (1). Interest in these organisms in terms of drug discovery is based on their treatment difficulty due to limited drugs and intrinsic resistance to current clinically used standard of care (SoC) drugs. Category A pathogens pose the highest risk to public health because they are rapidly progressing, difficult to treat and result in high mortality (2). Category B pathogens pose less risk because they are less transmittable with moderate morbidity rates and lower mortality (2). Novel drug candidates with unique pharmacological structures that circumvent existing drug resistance mechanisms and target previously unexploited molecular targets are urgently needed. Such candidates are envisioned to expand the structural diversity and increase the number of compounds progressing through the drug discovery pipeline for difficult to treat medically important pathogens. However, evaluation of drug candidates against NIAID Category A and B priority pathogens, emerging Gram-negative bacterial pathogens, and pre-IND enabling studies present several challenges. In particular, the spectrum of susceptibility, the infectious nature, and high mortality rates associated with these pathogens make drug discovery costly and resource intensive.

Evaluation of drug candidates begins with testing for potency, which is often performed against a reference laboratory-adapted bacterial strain or a closely related model organism (3-5). While laboratory or model strains provide a single reference for comparison of drug candidate performance, they do not adequately represent the spectrum of drug susceptibility of clinical strains (6). Analysis of the drug susceptibility profiles to SoC drugs against a spectrum of clinically derived strains in addition to reference laboratory strains can provide predictive information about a drug candidates performance in the clinic, but thus far strains with a broad range of susceptibilities encountered in clinical disease is not routinely incorporated early in drug activity evaluation. The variety of SoC drugs with activity, and their different modes of action makes it difficult to derive performance information from the susceptibility of a limited number of clinically used drugs and laboratory reference strains. Further, if new drug candidates are envisioned to replace or be employed in co-delivery regimens it is important to understand SoC drugs activity and mode of action. Accordingly, a panel of NIAID Category A and B priority and emerging strains used in the NIAID drug screening program has been assembled that includes reference and laboratory strains and strains that represent the naturally occurring susceptibilities of strains encountered during clinical infection (7). The species in this panel are *Burkholderia pseudomallei, Burkholderia mallei, Francisella tularensis, Yersinia pestis* and *Bacillus anthracis*.

Since there are no comprehensive reports of comparative susceptibility profiles of SoC drugs against this panel of strains that are routinely used in drug screening, we evaluated the susceptibility of the strains to 40 SoC drugs and determined susceptibility based on CLSI criteria (8). Susceptibility testing against a range of SoC drugs with various modes of action and unbiased hierarchical clustering analysis allowed us to: [1] establish a comparison resource of the clinical strains and reference laboratory strains for assessing the performance of drug candidates; [2] identified subsets of non-redundant strains that are able to classify the susceptibility range for each species; [3] discriminate pharmacophore classification for each species. This information can be used to better utilize limited resources and streamline drug candidate throughput in novel drug discovery or repurposing efforts to guide the selection of drug classes and to advance drug candidates with the greatest potential to treat emerging infectious diseases caused by these pathogens.

## Materials and Methods

### NIAID Bacterial strains for drug candidate assessment

The panel of Category A and B pathogens used for drug candidate evaluation in the NIAID drug screening program (7) consists of *Bacillus anthracis* (N=15), *Yersinia pestis* (N=5), *Francisella tularensis* (N=6), *Burkholderia mallei* (N=7), and *Burkholderia pseudomallei* (N=17) (Table 1). *Escherichia coli, Klebsiella pneumonia, Pseudomonas aeruginosa and Acinetobacter baumannii* are also included in the panel as quality and testing control strains. This strain panel is maintained as low passage stocks and stored at -80°C.

**Table 1.**
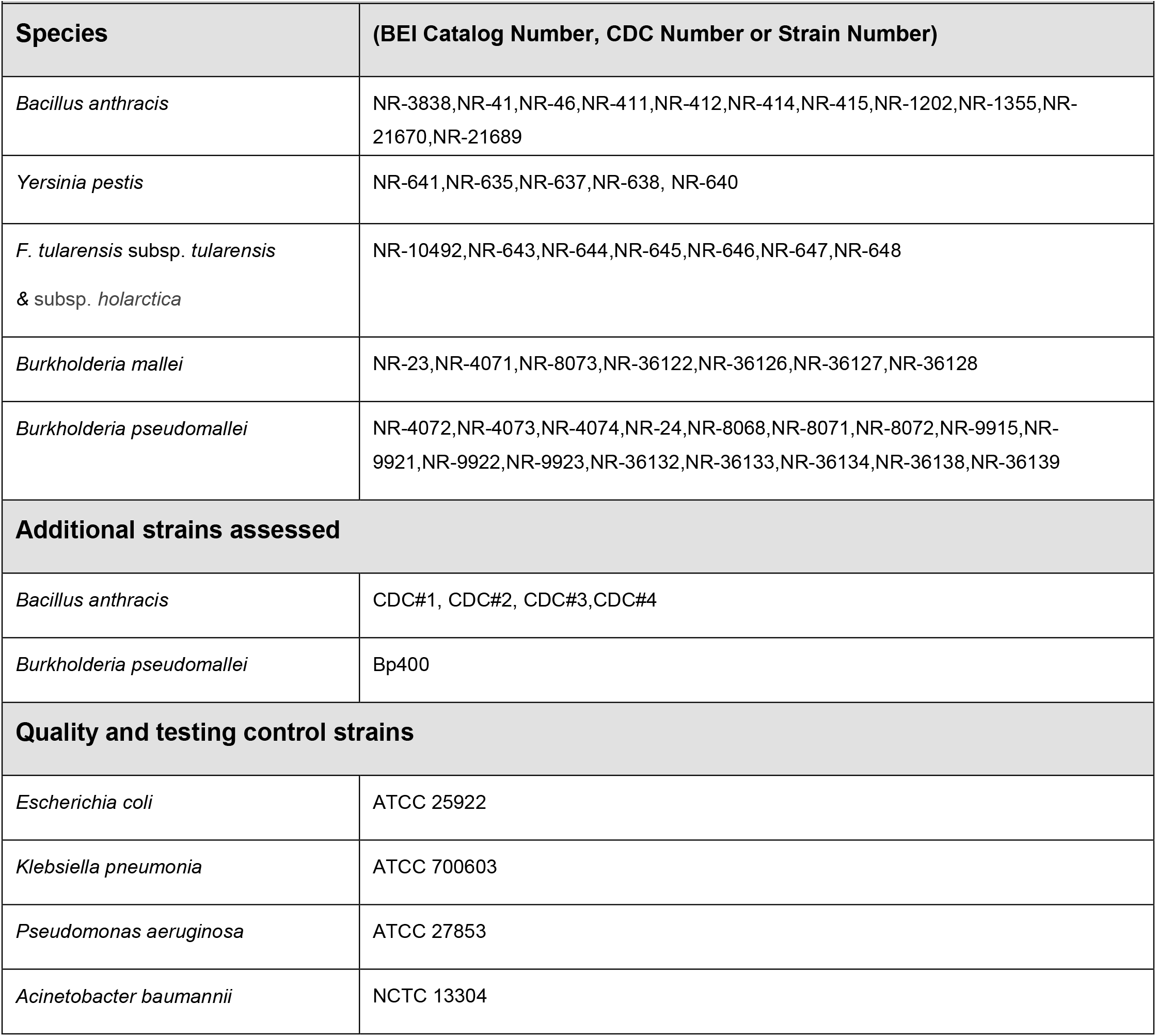
Category A pathogen strains

### Growth and maintenance conditions

#### B. anthracis

10ml Tryptic Soy Broth (TSB) (BD Franklin Lakes, NJ) was inoculated with 0.01 mL *B. anthracis* glycerol bacterial stock and incubated overnight at 37°C on an orbital shaker. Overnight cultures were diluted 1:100 into 10 mL TSB and incubated for an additional 6 hours. OD_600_ was taken and cultures were diluted to a concentration of 1⨯10^6^ CFU/mL in cation-adjusted Mueller-Hinton Broth (caMHB) (BD). 0.05mL of diluted cultures were used to inoculate 96-well plates.

#### Y. pestis

50mL Brain Heart Infusion Broth (BHI) (BD) was inoculated with 0.05mL *Y. pestis* glycerol bacterial stock and incubated for 48 hours at 28°C. OD_600_ was taken and cultures were diluted to a concentration of 1⨯10^6^ CFU/mL in cation-adjusted Mueller-Hinton Broth (caMHB) (BD). 0.05mL of diluted cultures were used to inoculate 96-well plates.

#### F. tularensis

*F. tularensis* was streaked onto Cystine Heart Agar supplemented with 2% Hemoglobin (CHAB) (BD) and incubated at 37°C for 72 hours. Bacterial suspensions were prepared in cation-adjusted Mueller-Hinton Broth supplemented with IsoVitalex (VWR Radnor, PA), 0.1% dextrose (Sigma Aldrich, St. Louis, MO), and 0.025% ferric pyrophosphate (Sigma Aldrich) (MMH) to an OD_600_ of ∼0.5. Suspensions were diluted in MMH to a concentration of 1⨯10^6^ CFU/mL and 0.05mL of diluted cultures were used to inoculate 96 well plates.

#### *B. pseudomallei* and *B. mallei*

10ml Luria-Bertani broth (LB) (BD) was inoculated with 0.01mL *B. pseudomallei* or *B. mallei* glycerol bacterial stock and incubated overnight at 37°C on an orbital shaker. Overnight cultures were diluted 1:100 into 10mL LB and incubated for an additional 6 hours. OD_600_ was taken and cultures were diluted to a concentration of 1⨯10^6^ CFU/mL in cation-adjusted Mueller-Hinton Broth (caMHB) (BD). 0.05mL of diluted cultures were used to inoculate 96-well plates.

### Standard of care drug plate preparation

The compounds in the 40 clinical SoC drug panel represent the different drug families: aminoglycosides, macrolides, beta-lactams, cephalosporins and quinolones. Drug master plates were prepared in triplicate with 40 SoC drugs, separately or in combination, dissolved in appropriate solvents to concentrations of 0.8mg/mL and 0.1mg/mL, and stored at -20 °C prior to testing. Master plates were used to prepare drug test plates at 64μg/mL and 8 μg/mL in 0.05ml caMHB. Once diluted 1:1 with inoculum, the final testing concentrations were 32μg/mL and 4 μg/mL, respectively. Ten wells per 96 well plate contained cAMHB only to serve as growth controls. All testing was performed per CLSI guidelines (8).

### Resazurin percent growth reduction determination

Inoculated 96-well plates were incubated for 18 hours at 37°C for *B. pseudomallei, B. mallei*, and *F. tularensis* and 24 hours at 37°C for *Y. pestis*. Resazurin sodium salt (Sigma Adlrich) was dissolved in PBS (Sigma Adlrich) at a concentration of 0.11mg/mL and sterile filtered through 0.2μm filter. 10μL was added to each well of the inoculated 96-well plate, and plates were incubated for an additional 4 hours at 37°C. Optical density was measured at 570nm and 600nm for each plate and percent growth reduction calculated using extinction coefficients and the following formula:

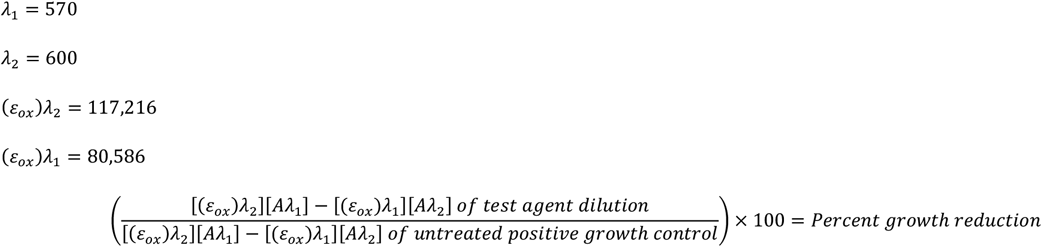

### Clustering and Subgroup identification

Manhattan distance and the sum of absolute difference were used to assess differences in the susceptibility profiles between subtypes within the five organisms utilized and between all strains combined. Utilizing the calculated Manhattan distances and the unweighted pair group method with arithmetic mean (UPGMA) clustering method allowed the identification of similarities between the different subtypes per species and for the combined subtypes of all species. Subtypes with distance equal to zero, indicating an exact match of their susceptibility profiles, were identified and replaced with one representative subtype. To obtain the minimum number of subtypes that span all possible drug susceptibilities after removing exact matching susceptibility profiles, we first identified the subtype that included the maximum number of drug susceptibilities. We then added the subtype that would, along with the first subtype, provide the maximum number of drug susceptibilities, given that that newly added subtype is either disjoint or not completely overlapping with the first. We continued adding subtypes utilizing the conditions described until adding a new subtype resulted in no new drug susceptibilities added to the spanning combined susceptibility profile.

## Results

### Source of clinical and laboratory strains and standard of care drugs

The bacterial strains tested are those used for drug candidate evaluation in the NIAID biodefense pathogens drug screening program (7). Additional strains included were *B. anthracis* strains CDC #1-4, and the comparator drug efflux deficient laboratory strain *B. pseudomallei* strain Bp400 (9, 10). In total, 50 strains including naturally occurring clinically associated *B. anthracis* (N=15), *Y. pestis* (N=5), *F. tularensis* (N=6), *B. pseudomallei* (N=17) and *B. mallei* (N=7) formed the strain panel were included in this study (Table 1).

The 15 *B. anthracis* strains include the standard lab strain Ames 3838, and the remaining strains were obtained from BEI resources or through the CDC. The 17 *B. pseudomallei* strains include the standard lab strains 1026b, K96243 and Bp400. The Bp400 strain is a double efflux knockout strain derived from 1026b and has been included in our drug screening program to determine the role of efflux in drug susceptibility (10). The seven *B. mallei* strains include the standard lab strain China7 (ATCC23344) and two strains from China7 lineages. The six *F. tularensis* strains include the standard lab strain SchuS4 and a second SchuS4 lineage strain. The five *Y. pestis* strains include the standard lab strain CO92. *E. coli* (ATCC25922), *K. pneumonia* (ATCC700603), *P. aeruginosa* (ATCC27853), and *A. baumannii* (NCTC13304) were used as quality and control strains.

Susceptibility of each bacterium to 40 SoC drug treatments, consisting of single drugs (n=34) and drug combinations (n=6), were determined by testing for percent growth inhibition consistent with CLSI guidelines (8). The compounds evaluated included drugs in the aminoglycoside, macrolide, penicillin, sulfa, carbapenem, cephalosporin and quinolone drug classes with known modes of action that target protein synthesis, cell wall synthesis, DNA replication, DNA synthesis and RNA synthesis. The MIC value for each strain was used to categorize the susceptibility of each strain to each drug or drug combination as susceptible, intermediate, or resistant. The cutoff determinations for resistance, intermediate, and susceptible were chosen based on CLSI guidelines and standard practices required for pre-IND filing and an FDA Type A, B or C meeting (8) (Table 2).

**Table 2.** Susceptibility Profile Cutoffs

| Table 2 – Susceptibility Profile Cutoffs |  |  |  |
| --- | --- | --- | --- |
| Cutoff Values µg/ml | Susceptible | Intermediate | Resistant |
| Single Compounds | <4 | 4 - 32 | >32 |
| Imipenem/cilastatin (1:1) | <4/4 | 4/4 - 32/32 | >32/32 |
| Amoxicillin/clavulanic Acid (4:1) | <4/1 | 4/1 - 32/8 | >32/8 |
| Piperacillin/tazobactam (8:1) | <4/0.5 | 4/0.5 - 32/4 | >32/4 |
| Ampicillin/sulbactam (2:1) | <4/2 | 4/2 - 32/16 | >32/16 |
| Sulfadiazine/pyrimethamine (1:20) | <0.25/5 | 0.25/5 - 2/40 | >2/40 |
| Trimethoprim/sulfamethoxazole (1:19) | <0.25/4.75 | 0.25/4.75 - 2/38 | >2/38 |

### Susceptibility to standard of care drugs and identification of minimal strain set

In order to determine the spectrum of strain susceptibility for each species to SoC drugs and to identify non-redundant strains that are representative of the clinical susceptibilities for each species, each strain was tested and classified as susceptible, intermediate or resistant based on CLSI guidelines (8) and subjected to an unbiased cluster analysis. Defined susceptibility profiles for each strain provides a comparative reference useful for determining the performance of drug candidates. Unbiased cluster analysis using strain susceptibility profiles groups strains based on similarity of susceptibility to the 40 SoC drug conditions tested and identifies strains with susceptibility profiles that were an exact match. The susceptibility values and unbiased analysis provided clear demarcations for strain grouping based on pathogen and drug susceptibility, which successfully clustered strains revealing the spectrum of susceptibility and exact match susceptibility profiles (Table 3). This resulted in a subset of non-redundant strains that represent the spectrum of clinical susceptibilities of each species tested, and when combined resulted in a non-redundant panel representing all the species.

**Table 3:** List of exact matches

| Table 3: List of exact matches |  |  |  |  |
| --- | --- | --- | --- | --- |
| Exact<br>Matches 1 | *B. anthracis*.Graves | *B. anthracis* Ames A0462 | *B. anthracis* A0471 |  |
| Exact<br>Matches 2 | *B. anthracis* 46 PY 5 | *B. anthracis* Kruger BA0442 | *B. anthracis* Vollum | *B. anthracis* A0318 |
| Exact<br>Matches 3 | *B. mallei*.China7 ATCC23344 | *B. mallei* China7 NBL 7 |  |  |
| Exact<br>Matches 4 | *B. pseudomallei* K96243 | *B. pseudomallei* MSHR465a |  |  |

#### F. tularensis susceptibility and identification of predictive strains

The *F. tularensis* strains were clustered by susceptibility profiles resulting in two distinct susceptibility groups. Susceptibility Group 1 contained strains WY96 and MA00 and included the most susceptible strains, with WY96 being susceptible to every treatment but bacitracin and MA00 being resistant to bacitracin, colistin, amoxicillin and carbenicillin, and intermediately susceptible to daptomycin, ampicillin, polymyxin and the combination of trimethoprim/sulfamethoxizole. Susceptibility Group 2 organisms demonstrated resistance or intermediate susceptibilities to approximately 50% of the SoC drugs. Strain SchuS4 FSC237 was the most resistant *F. tularensis* strain tested as it demonstrated resistance to 20 SoC treatments and intermediate susceptibility to an additional 2 SoC treatments. Similarly, SchuS4 was resistant to 13 SoC treatments and showed intermediate susceptibility to 9 SoC treatments. Strains KY99 and OR96 were resistant to 11 SoC treatments and intermediately susceptible to 9 and 7 SoC treatments, respectively (Figure 1A). A secondary analysis was performed to determine the minimal *F. tularensis* strain panel that represented the resistance profiles of the entire panel, which identified the 4 strains MA00, SchuS4 FSC237, SchuS4 and KY99 (Figure 1B). No strains were identified that were an exact match for drug susceptibility.

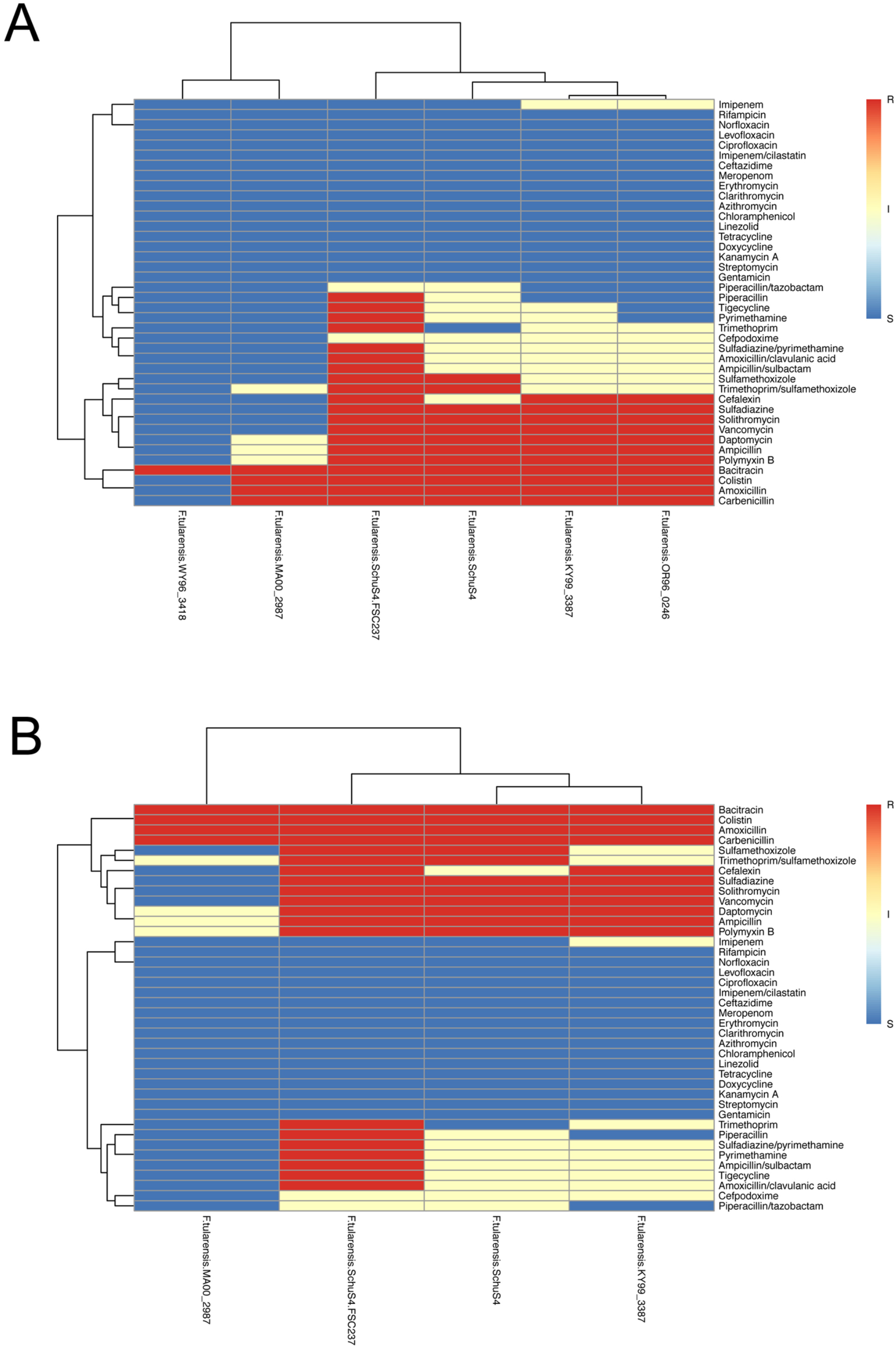

#### B. anthracis susceptibility and identification of non-redundant strains

The 15 strains of *B. anthracis* were grouped by susceptibility profiles resulting in identification of a singleton strain and 2 susceptibility groups (Figure 2A). *B. anthracis* strain CDC #1 (20000032823) was the singleton strain with the furthest neighbor at an R^2^ value of 0.69. *B. anthracis* susceptibility Group 2 (R^2^ = 0.97) included strains 525, 506, Ames 3838, CDC #3 (2010719149), CDC #2 (2002734753), and CDC #4 (20060200760). Susceptibility Group 3 (R^2^ = 0.98) contained *B. anthracis* strains WNA, Graves, Ames A0462, A0471, 46-PY-5, Kruger B, Vollum, and A0318 (Figure 2A). The susceptibility profile of the singleton strain CDC#1 is similar to the susceptibilities of Group 2, except strain CDC#1 is resistant to the beta-lactams amoxicillin, ampicillin, carbenicillin and piperacillin. The susceptibilities that distinguish susceptibility Groups 2 and 3 are those to solithomycin, cephalexin and the ampicillin/sulbactam combination. Susceptibility Group 2 strains are susceptible to these beta-lactams, while few of the strains assigned to Susceptibility Group 3 have intermediate susceptibilities to piperacillin. Secondary analysis revealed exact matches in drug susceptibility for *B. anthracis* strains Graves, Ames A0462, A0471, and exact matches in drug susceptibility for strains 46-PY-5, Kruger B, Vollum, and A0318 (Table 3). Together, this identified *B. anthracis* strains CDC#1, WNA, Graves, 525, 506 and CDC#2 as representatives of the entire strain panel (Figure 2B).

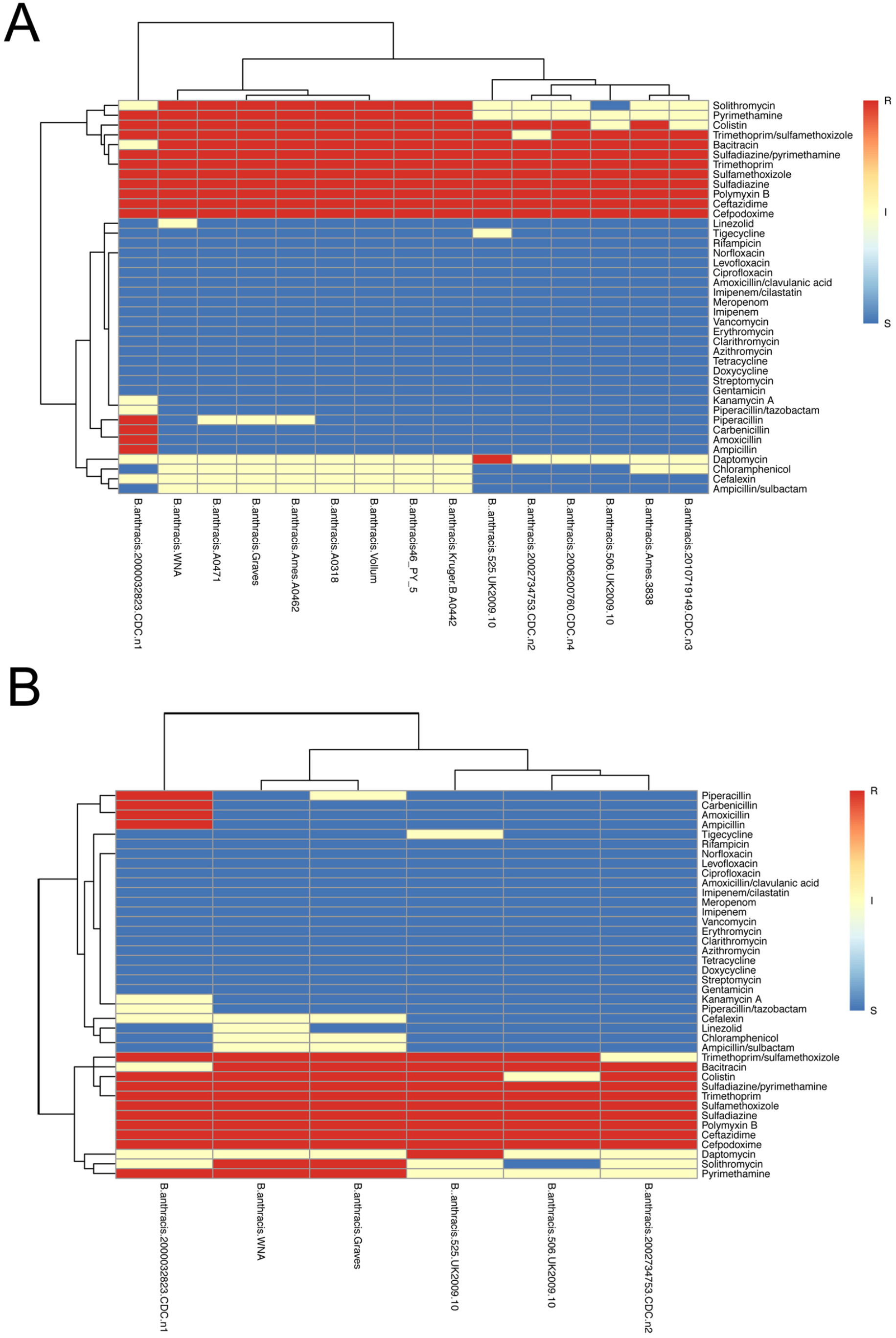

#### Y. pestis strains and identification of non-redundant strains

The 5 *Y. pestis* strains were grouped by susceptibility profiles, resulting in identification of a singleton strain and 1 susceptibility group. The singleton strain was Nepal516 (R^2^ = 0.74), and the PEXU2, PB6, ZE94, and CO92 strains clustered in the susceptibility group (Figure 3A). A secondary analysis was performed to determine the minimal strain panel, which only eliminated the *Y. pestis* strain PB6 as a necessary strain to represent clinical susceptibility of the strain panel (Figure 3B). No strains were identified that were an exact match for drug susceptibility.

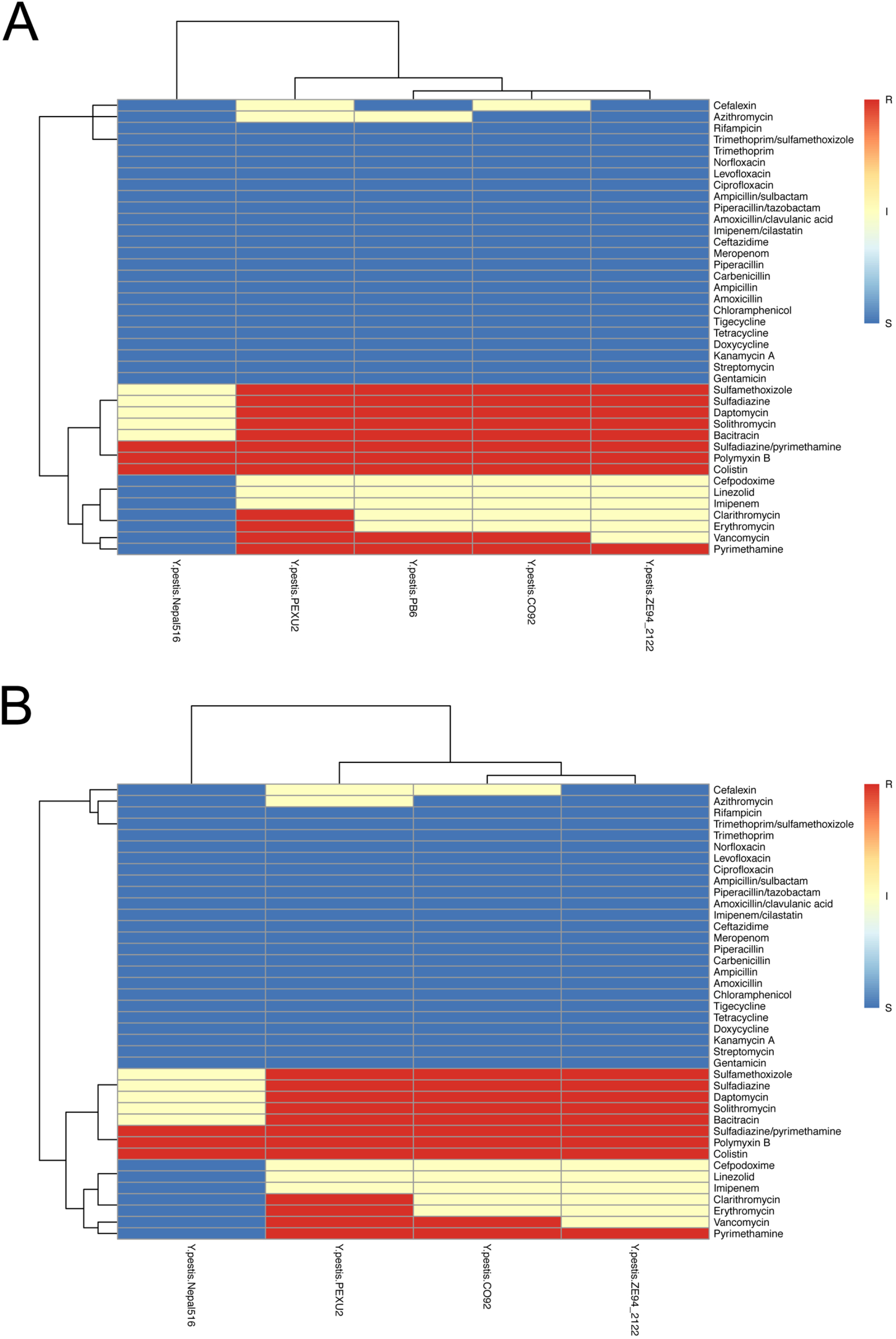

#### B. pseudomallei susceptibility and identification of non-redundant strains

The susceptibility profile of 17 strains of *B. pseudomallei* were analyzed, which revealed three furthest neighbor singletons, DD503 (R^2^ = 0.77), MSHR435 (R^2^ = 0.85), and NCTC6700 (R^2^ = 0.90), and several susceptibility groups (Figure 4A). Although *B. pseudomallei* Bp400 distributed as a singleton (R^2^ = 0.57), it should not be considered a clinical variant since it is an efflux deficient tool strain derived from 1026b that significantly alters its susceptibility profile. Susceptibility Group 1 (R^2^ = 0.92) contained strains 1710b, 406e, 1106b, and NCTC7383. Susceptibility Group 2 (R^2^ = 0.94) included strains China3, 1026b, and 1710a. Susceptibility Group 3 (R^2^ = 0.94) contained strains MSHR668, K96243, MSHR465a, NCTC7431, NCTC10274, and NCTC10276. *B. pseudomallei* DD503 is the most susceptible clinical strain with resistance limited to vancomycin, amoxicillin, streptomycin, macrolides and polypeptides. The singleton strains NCTC6700 and MSHR435 are the most resistant *B. pseudomallei* strains. Strain NCTC6700 is only completely susceptible to imipenem, doxycycline, tetracycline, tigecycline, ciprofloxacin and levofloxacin. Strain MSHR435 is susceptible to imipenem, ciprofloxacin and levofloxacin, but resistant to doxycycline, tetracycline, tigecycline. Susceptibility Groups 1-3 have intermediate susceptibility profiles marked by intermediate susceptibilities to chloramphenicol, ceftazidime, gentamicin, kanamycin, norfloxacin, sulfadiazine, ampicillin, sulfamethoxozole and combinations. A secondary analysis was performed to determine the minimal strain panel that represents the clinical susceptibilities of these 17 testing strains. This analysis revealed exact matches in drug susceptibility for *B. pseudomallei* strains K96243 and MSHR465a (Table 3) and identified strains MSHR435, NCTC7383, 1026b, NCTC6700, Bp400, and DD503 as representatives of the strain panel (Figure 4B). *B. pseudomallei* Bp400 is also useful as a strain in the non-redundant test panel for comparison to 1026b to discern the impact of drug efflux on the observed susceptibility.

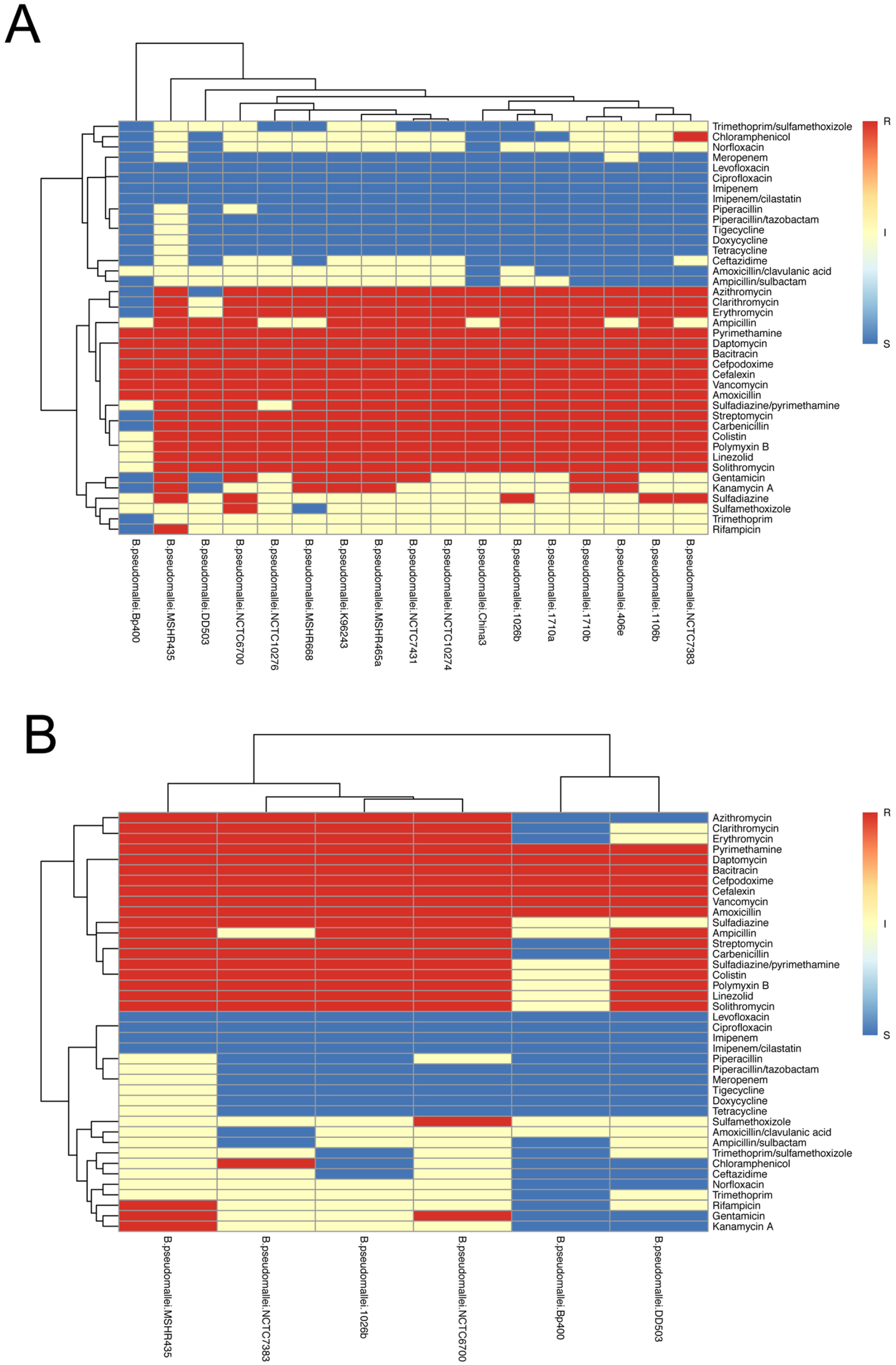

#### B. mallei susceptibility and identification of non-redundant strains

The susceptibility profile of 7 strains of *B. mallei* were clustered, and strains 10248 (R^2^ = 0.76), NCTC10260 (R^2^ = 0.86) and NCTC120 (R^2^ = 0.90) were identified as singletons, while strains GB8 Hourse4, NCTC12938, China7 ATCC23344, and China7 NBL7 were organized into Susceptibility Group 1 (R^2^ = 0.98) (Figure 5A). This organization resulted from resistance and intermediate susceptibilities to the cephalosporins, cephalexin and cefpodoxime, the beta-lactams, ampicillin, amoxicillin and carbenicillin, as well as colistin, polymyxin, bacitracin and daptomycin. Secondary analysis revealed that the susceptibilities of *B. mallei* strains China7 NBL7 and China7 ATCC23344 were exact matches (Table 3) determining that the clinical susceptibilities of these 7 testing strains could be represented by the *B. pseudomallei* strains GB8 Horse4, 10248, NCTC10260 and NCTC120 (Figure 5B).

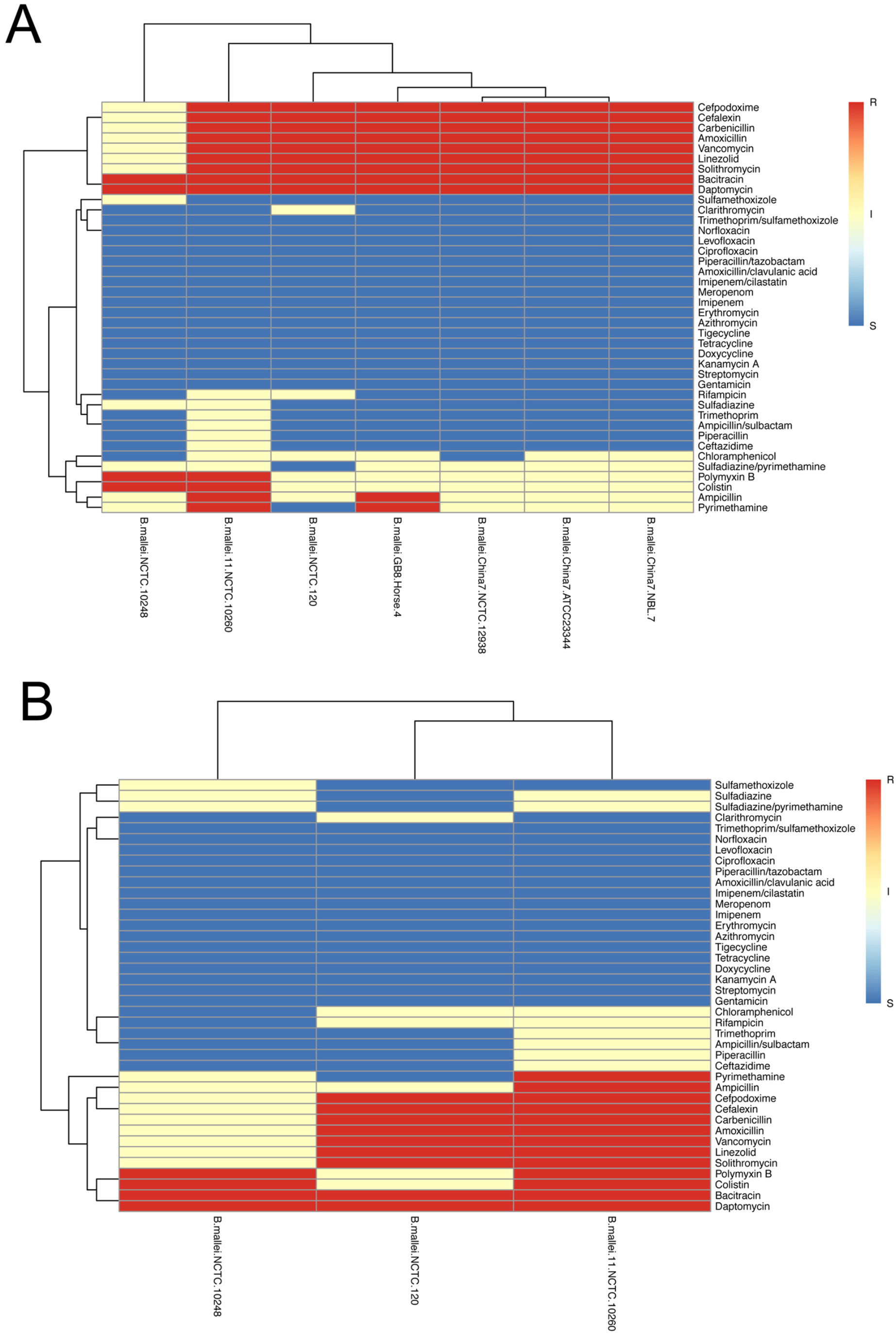

### Identification of minimal non-redundant strain set for Category A and B drug testing

Testing exploratory and lead drug candidates against 50 category A and B strains from 5 different bacterial species requires significant resources. The screening efficiency can be much improved using an initial pan-species screening panels of strains representative of the full spectrum drug susceptibilities and maximum resistance profiles. Drug candidates that exceed screening criteria in this initial screen can then be progressed to testing against larger panels of strains if confirmation is necessary. In order to identify representative strains from each pathogen species, an unbiased cluster analysis was performed with all 50 of the strains to categorize them based on susceptibility profile and to identify and eliminate strains from each species that had susceptibility profiles that were an exact match. This unbiased analysis successfully clustered strains into 3 distinct groups (Figure 6A). Group 1 contains only *B. pseudomallei* strains, group 2 consists of subgroups containing *Y. pestis, B. mallei, B. pseudomallei* and *F. tularensis* strains and group 3 contains only *B. anthracis* strains. Removal of strains with exact matches (Table 3) resulted in a reduced pan-species panel consisting of 9 strains representing all 5 species based on the spectrum of susceptibilities to SoC drugs tested (Figure 6B). These 9 strains consisted of 3 *B. pseudomallei*, 2 *B. mallei*, 1 *Y. pestis*, 1 *F. tularensis* and 2 *B. anthracis* strains. Further, analysis was performed to identify a single strain from each species that is most resistant to the SoC drugs tested, which provides a focused pan-species screening panel of maximum resistance. The strains in this screening panel are *F. tularensis* SchuS4-FSC237, *B. anthracis* CDC#1, *Y. pestis* PEXU2, *B*.*pseudomallei* MSHR435 and *B. mallei* 10260. This substantiated that individual strain susceptibility profiles to SoC drugs could accurately classify strains into non-redundant groups based on susceptibility profiles that can be used in pan-species screening panels for initial drug screening strategies.

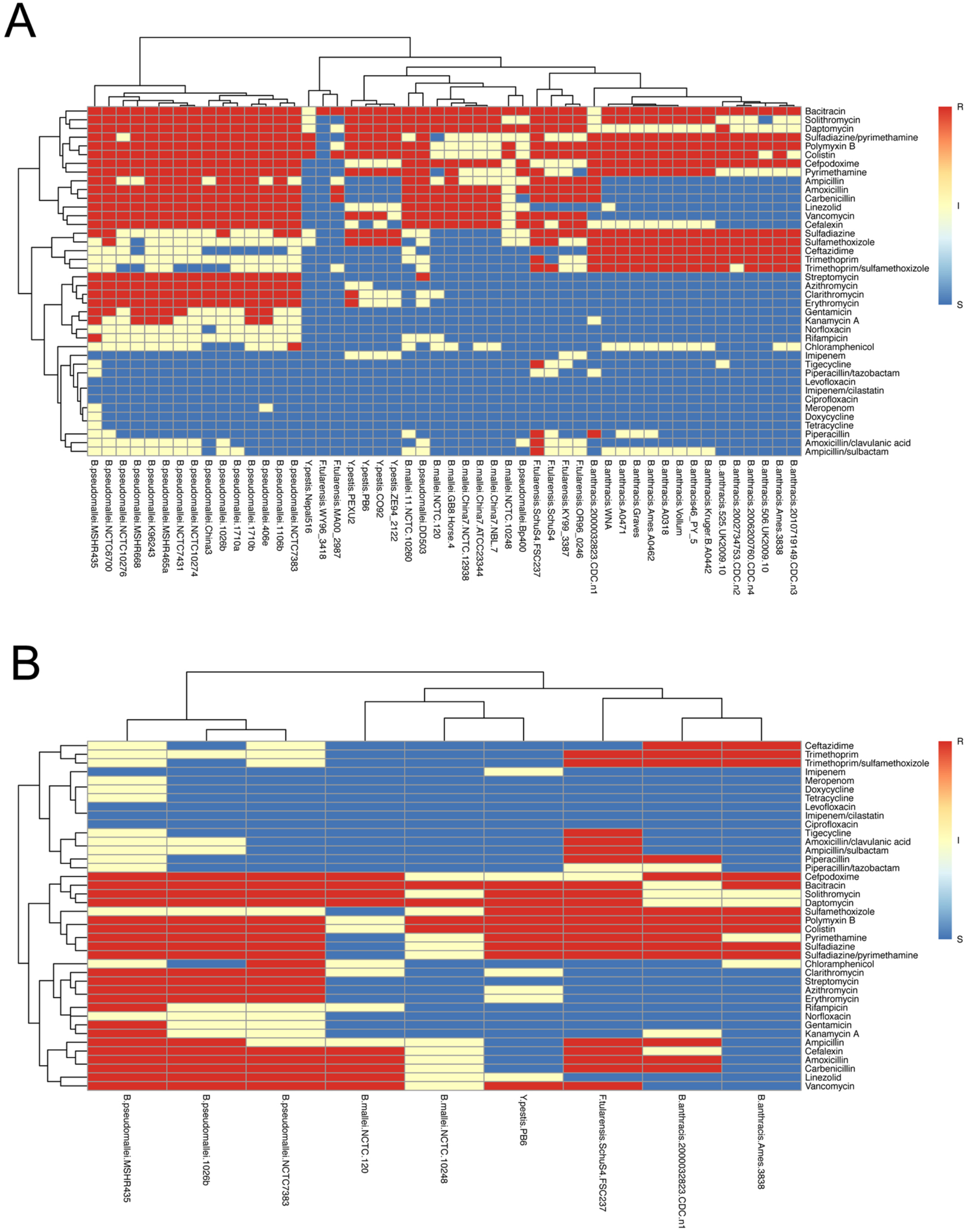

### Pharmacophore classification for Category A and B pathogens

Heterogeneity in drug resistance was observed across the total number of strains for each species, indicating a broad-spectrum of susceptibilities observed in clinically obtained strains in each species. General resistance for all the strains within *B. anthracis, B. pseudomallei, B. mallei, F. tularensis and Y. pestis* was observed for polymyxin, polypeptide, lipodpetide drug classes cephalosporin and penicillin, and to a lesser degree macrolide antibiotics (Table 4). Another important finding was the standard laboratory strains routinely used in drug screening programs do not accurately approximate the susceptibility of the most resistant clinical strains in any of the species. Another important finding was each species is uniquely susceptible to specific drug classes.

**Table 4.** Percent drug resistance of bacterial strains

| Class | Drug | % of Resistant Strains |  |  |  |  |
| --- | --- | --- | --- | --- | --- | --- |
|  |  | Ba | Bp | Bm | Ft | Yp |
| Protein Synthesis Inhibitors | Streptomycin | 0% | 94% | 0% | 0% | 0% |
| Protein Synthesis Inhibitors | Gentamicin | 0% | 47% | 0% | 0% | 0% |
| Protein Synthesis Inhibitors | Kanamycin A | 0% | 35% | 0% | 0% | 0% |
| Protein Synthesis Inhibitors | Doxycycline | 0% | 0% | 0% | 0% | 0% |
| Protein Synthesis Inhibitors | Tetracycline | 0% | 0% | 0% | 0% | 0% |
| Protein Synthesis Inhibitors | Tigecycline | 0% | 0% | 0% | 17% | 0% |
| Protein Synthesis Inhibitors | Linezolid | 0% | 94% | 86% | 0% | 0% |
| Protein Synthesis Inhibitors | Chloramphenicol | 0% | 6% | 0% | 0% | 0% |
| Protein Synthesis Inhibitors | Azithromycin | 0% | 88% | 0% | 0% | 0% |
| Protein Synthesis Inhibitors | Clarithromycin | 0% | 88% | 0% | 0% | 20% |
| Protein Synthesis Inhibitors | Solithromycin | 53% | 94% | 86% | 67% | 80% |
| Protein Synthesis Inhibitors | Erythromycin | 0% | 88% | 0% | 0% | 20% |
| Cell Wall Synthesis Inhibitors | Vancomycin | 0% | 100% | 86% | 67% | 60% |
| Cell Wall Synthesis Inhibitors | Amoxicillin | 7% | 100% | 86% | 83% | 0% |
| Cell Wall Synthesis Inhibitors | Ampicillin | 7% | 65% | 29% | 67% | 0% |
| Cell Wall Synthesis Inhibitors | Carbenicillin | 7% | 94% | 86% | 83% | 0% |
| Cell Wall Synthesis Inhibitors | Piperacillin | 7% | 0% | 0% | 17% | 0% |
| Cell Wall Synthesis Inhibitors | Imipenem | 0% | 0% | 0% | 0% | 0% |
| Cell Wall Synthesis Inhibitors | Meropenem | 0% | 0% | 0% | 0% | 0% |
| Cell Wall Synthesis Inhibitors | Cefalexin | 0% | 100% | 86% | 50% | 0% |
| Cell Wall Synthesis Inhibitors | Ceftazidime | 100% | 0% | 0% | 0% | 0% |
| Cell Wall Synthesis Inhibitors | Cefpodoxime | 100% | 100% | 86% | 0% | 0% |
| Cell Wall Synthesis Inhibitors | Imipenem/cilastatin | 0% | 0% | 0% | 0% | 0% |
| Cell Wall Synthesis Inhibitors | Amoxicillin/clavulanic acid | 0% | 0% | 0% | 17% | 0% |
| Cell Wall Synthesis Inhibitors | Piperacillin/tazobactam | 0% | 0% | 0% | 0% | 0% |
| Cell Wall Synthesis Inhibitors | Ampicillin/sulbactam | 0% | 0% | 0% | 17% | 0% |
| Cell Wall Synthesis Inhibitors | Polymyxin B | 100% | 94% | 29% | 67% | 100% |
| Cell Wall Synthesis Inhibitors | Colistin | 87% | 94% | 29% | 83% | 100% |
| Cell Wall Synthesis Inhibitors | Bacitracin | 93% | 100% | 100% | 100% | 80% |
| Cell Membrane Function Inhibitors | Daptomycin | 7% | 100% | 100% | 67% | 80% |
| DNA Replication Inhibitors | Ciprofloxacin | 0% | 0% | 0% | 0% | 0% |
| DNA Replication Inhibitors | Levofloxacin | 0% | 0% | 0% | 0% | 0% |
| DNA Replication Inhibitors | Norfloxacin | 0% | 0% | 0% | 0% | 0% |
| DNA Synthesis Inhibitors | Sulfadiazine | 100% | 29% | 0% | 67% | 80% |
| DNA Synthesis Inhibitors | Sulfamethoxazole | 100% | 6% | 0% | 33% | 80% |
| DNA Synthesis Inhibitors | Pyrimethamine | 60% | 100% | 29% | 17% | 80% |
| DNA Synthesis Inhibitors | Trimethoprim | 100% | 0% | 0% | 17% | 0% |
| DNA Synthesis Inhibitors | Sulfadiazine/pyrimethamine | 100% | 88% | 0% | 17% | 100% |
| DNA Synthesis Inhibitors | Trimethoprim/sulfamethoxazole | 93% | 0% | 0% | 33% | 0% |
| RNA Synthesis Inhibitors | Rifampicin | 0% | 6% | 0% | 0% | 0% |

Ba: *Bacillus anthracis*; Bp: *Burkholderia pseudomallei*; Bm: *Burkholderia mallei*; Ft: *Francisella tularensis*; Yp: *Yersinia pestis*

*B. anthracis* strains were most susceptible to ansamycins, carbapenem/β-lactamase inhibitors, carbapenems alone, glycopeptides, tetracyclines, and aminoglycosides (Table S1). All the *F. tularensis* strains were susceptible to 17 of the SoC drugs, and the majority of the strains were susceptible to drugs in the quinolone, macrolide, carbapenem and aminoglycoside, and tetracycline drug classes, and the addition of a DHFR inhibitor to DHPS inhibitors increased the susceptibility of *F. tularensis*. (Table S2). The *Y. pestis* strains in the panel were fully susceptible to aminoglycosides, tetracycline, penicillin and cephalosporin antibiotics (Table S3). *B. pseudomallei* was the most resistant species with intermediate susceptibility to ansamycin, the DHFR/DHPS combination, DHFR alone, sulfonamide, and amphenicol drugs. The *B. pseudomallei* drug efflux deficient comparator strain Bp400 was only resistant to vancomycin, amoxicillin, bacitracin, daptomycin and pyrimethamine, which indicates that the observed resistance to these drugs in wild-type strains of *B. pseudomallei* are not due to efflux. This observation is consistent with previous reports and the well-known drug efflux mechanisms that are encoded by *B. pseudomallei* (9-11). The four drug groups that showed potency against *B. pseudomallei* are quinolones, tetracyclines, carbapenem, and penicillin/β-lactamase combinations (Table S4). In general, the related *B. mallei* strains were more susceptible to the SoC compounds tested than *B. pseudomallei*, which is consistent with the reduced drug efflux capabilities compared to *B. pseudomallei* (Table S5). Notably, doxycycline, tetracycline and tigecycline, imipenem, piperacillin and meropenem, ciprofloxacin, levofloxacin and norfloxacin are the only SoC drugs were widespread resistance was not observed for any of the strains across the 5 species tested.

## Discussion

Assessment of drug candidates relies on multi-step activity screening strategies where compounds are typically assessed against a laboratory reference strain or model strain. Drug candidates that demonstrate activity are then screened against additional bacterial strain panels, preferably including clinical strains or strains that fully represent the susceptibility range of clinical strains when known. It is often assumed that the additional strains, particularly strains obtained from clinical settings, fully represent the extent of drug susceptibility present in contemporary clinical strains causing infections. While reference strains are useful for determining a drug candidates’ “general” activity, they are most often laboratory-adapted strains, and therefore cannot be assumed to completely reflect the true drug susceptibilities of the strains encountered in the clinical setting. Unless laboratory and clinical strains with known activities for a large number of SoC drugs are included in screening panels, it is difficult to interpret the performance of drug candidates and benchmark their potential for clinical use, particularly for broad spectrum candidates. Our evaluation of these Category A and B priority strains including strains of clinical origin as well as the reference laboratory strains revealed stark differences in the drug profiles between laboratory strains and clinical strains, and considerable heterogeneity in drug susceptibly among the clinical strains. These finding support the utility of evaluating active drug candidates using non-redundant primary screening panels or pan-species screening panels followed by evaluation in secondary strain panels that include additional clinical strains for confirmation of activity. We also found that the clinical susceptibly spectrum for each of the Category A and B priority pathogen strain panels can be represented by fewer strains than originally selected for screening panels. In several cases the drug susceptibility profiles of several strains were exact matches, and therefore the redundant strains provided no additional screening value. This study clearly defined species and pan-species drug screening panels consisting of the strains routinely used to evaluate the performance of drug candidates (Figure 7). However, the combinations of strains in each panel are different than those currently used in screening programs.

**Figure 7.**
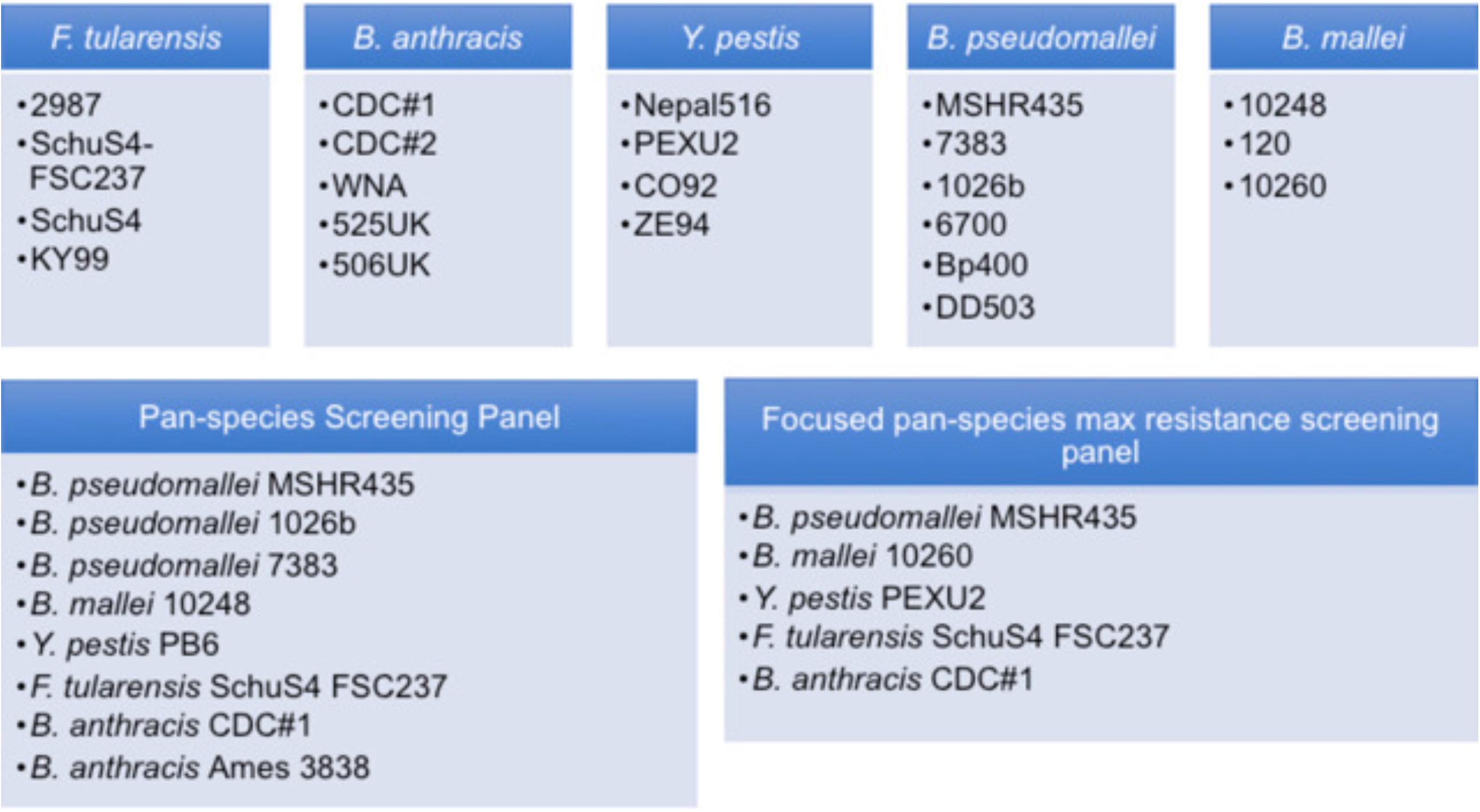
Non-redundant Screening Panels.

Visualization of susceptibility patterns for strains in each species also revealed susceptibility-resistant patterns for drug classes, and drug classes with generalized broad-spectrum activity. Notably, it was found that all Category A and B species tested were generally susceptible to doxycycline, tigecycline, tetracycline, chloramphenicol, piperacillin, imipenem, meropenem, rifampicin, norfloxicin, ciprofloxacin, levofloxacin, and piperacillin/tazobactam, imipenem/cilastatin, amoxicillin/clavulanic acid and ampicillin/sulbactam combinations making these SoC treatments a good first choice for broad-spectrum treatment for infections from an unknown agent. Most species were susceptible to Streptomycin, gentamicin, Kanamycin A, azithromycin, clarithromycin, solithromycin, erythromycin, and ceftazidime making these SoC drugs a good second choice or for combination treatment. *B. anthracis* was uniquely more susceptible to ampicillin, amoxicillin, carbenicillin and vancomycin compared to the other priority pathogens. *F. tularensis, B. mallei* and *Y. pestis* species were more susceptible to sulfadiazine, sulfamethoxazole, ceftazidime, and trimethoprim. *B. pseudomallei* was found to be the most naturally resistant species not being uniquely sensitive to any of the SoC drug tested and generally more resistant to chloramphenicol, clarithromycin, streptomycin azithromycin, erythromycin, ampicillin, amoxicillin, carbenicillin and vancomycin. Even the combination of beta-lactamase inhibitors, clavulanic acid and sulbactam, only slightly improved susceptibility, further substantiating the role of drug efflux in the general drug resistance of *B. pseudomallei* (9, 11). The majority of the strains in this study were susceptible to all the drugs in the tetracycline and quinolone drug classes and resistant to macrolide and penicillin class. Notably, the penicillin drugs only showed activity when paired with beta-lactamase inhibitor.

Classification of bacterial strains and species based on susceptibility profiles to SoC drugs can be used to guide strain selection for drug screening efforts and provide information about active drug classes and general drug resistance patterns. Susceptibility profiles of a large number of SoC drugs representing a variety of pharmacophores can also be exploited for rational drug design and drug repurposing efforts. In both cases, this information can afford improved utilization of limited resources and streamline drug candidate throughput. This information can also be used to guide treatment strategies for single and drug combination regimens that can be used to treat an infection when the agent is known or even before the specific pathogen is confirmed by laboratory testing.

## Acknowledgement

We gratefully acknowledge Julie Starkey for assisting with editing and for scientific input. This work was supported by funds from the Department of Microbiology, Immunology and Pathology Strategic Initiative at Colorado State University (R.A.S.), NIAID Task Order A30 (R. A. S.).

## Author contributions

The determination of MIC was performed by J. E. C., data structural analysis by Z.A. Study design and writing was performed by J. E. C., Z.A. and R. A. S..

